# SPONTANEOUS VISUAL IMAGERY DURING EXTENDED MUSIC LISTENING IS ASSOCIATED WITH RELIABLE ALPHA SUPPRESSION

**DOI:** 10.1101/2025.05.18.654076

**Authors:** Sarah Hashim, Diana Omigie

**Affiliations:** Department of Psychology, Goldsmiths, University of London, United Kingdom

**Author notes:** Corresponding author: Sarah Hashim, Department of Psychology Goldsmiths, University of London 8 Lewisham Way, New Cross, London, UK SE14 6NW, SH.

**Keywords:** music listening, EEG, visual imagery, spontaneous, deliberate, intentionality, probe-caught methodology

## Abstract

Music is widely recognised as being able to evoke images in the mind’s eye. However, the neural basis of visual imagery experiences during music listening remains poorly understood. Here, we combined probe-caught experience sampling methodology with 32-channel electroencephalography (EEG) recordings in order to investigate the neuro-oscillatory correlates of music-evoked visual imagery and examine how spontaneously generated imagery compares to more deliberately generated forms. Thirty participants listened with closed eyes to four blocks of music, differing in their familiarity and relaxation potential and spanning a range of genres. In response to probes sent throughout listening, participants indicated whether or not they had been experiencing visual imagery and, if they had, whether the experienced visual imagery had been spontaneous or deliberate. Cluster permutation analyses on the time-frequency decomposed EEG data revealed alpha power suppression during visual imagery that was more reliable during spontaneous than deliberate imagery. Further, while theta and delta bands did not discriminate the presence or absence of the visual imagery experience or its intentionality subtypes, we observed that gamma power suppression in fronto-central areas was present during visual imagery experiences. Our results extend prior findings of a role of posterior alpha suppression in visual imagery to show its reliability in music-evoked spontaneous imagery specifically. We consider plausible interpretations of the presence and absence of other oscillatory signatures in relation to the listening conditions used in the current study.

**Highlights:**

- Probe-caught method shown to be effective in capturing moments of music-evoked visual imagery
- Evidence for a high prevalence of spontaneous music-evoked imagery
- Spontaneous music-evoked visual imagery shown to be associated with posterior alpha suppression

## 1. INTRODUCTION

The internalisation of one’s attention is a common mental action that leads to a variety of overlapping states including visual mental imagery (Pearson, 2019), mind wandering (Christoff et al., 2016; Martarelli et al., 2016), and creative thinking (Belfi et al., 2017; Luft et al., 2019). Music listening, an activity often used as an escape from the outside world (Taruffi et al., 2017), has a widely recognised ability to evoke visual imagery in listeners, with some studies suggesting music-evoked visual imagery occurs in at least 70% of listeners (Küssner & Eerola, 2019; Vuoskoski & Eerola, 2015). Recent research has begun to characterise the different forms that music-evoked visual imagery can take (e.g., Dahl et al., 2022; Groves et al., 2023; Hashim et al., 2023; Küssner & Eerola, 2019) and to reveal how these different forms are reflected in brain activity (e.g., Fachner et al., 2019; Hashim et al., 2024). However, few studies into music-evoked visual imagery have specifically addressed the question of how often such experiences happen spontaneously as opposed to with deliberate intention. Furthermore, and more pertinently here, no study has examined the extent to which spontaneous and deliberate forms of music-evoked visual imagery implicate differing patterns of neural activity.

In mind wandering research, authors have proposed the existence of two attentional modes of mind wandering, namely intentional mind wandering and unintentional mind wandering (Seli, Risko, & Smilek, 2016; Seli, Risko, Smilek, et al., 2016). While the intentionality of visual imagery has not yet been a focus of research, the literature nevertheless suggests that there is a basis for distinguishing visual imagery that occurs spontaneously from that occurring with deliberate intention (Taruffi & Küssner, 2019). Specifically, alongside studies that have examined and reported on deliberate forms of imagery following explicit instruction (Cespedes-Guevara & Dibben, 2022; Day & Thompson, 2019; Hashim et al., 2020, 2023; Herff et al., 2022; Küssner & Eerola, 2019), other studies have examined and reported on more spontaneous forms of visual imagery, as might occur, for example, during deep meditation (Luft et al., 2019).

### 1.1. Capturing the occurrence of music-evoked visual imagery

In one of the few studies to date on the neural correlates of music-evoked visual imagery (Hashim et al., 2024), participants’ EEG was recorded while they listened to musical excerpts and rated the extent to which they experienced static and dynamic imagery during said excerpts. That study was important in confirming that qualities of music-evoked visual imagery – like the presence of motion – can be seen reflected in neural data. However, by requiring participants to report on how much imagery they experienced on each trial, that study arguably encouraged deliberate, rather than more spontaneous forms of imagery, and as such it remains an open question how relevant the signatures reported there are to spontaneously generated imagery. In any case, by not directly inquiring as to whether any visual imagery that was experienced was spontaneously or deliberately generated, that and other previous studies have been unable to speak to how the neural correlates of these visual imagery types may differ.

A large body of mind wandering research has showcased the usefulness of the probe-caught methodology (Smallwood & Schooler, 2015) for capturing nuanced differences in those moments when individuals shift their attention inward. However, the probe caught experience method is yet to be fully exploited in music listening studies. One study by Taruffi and colleagues (2017) used a simple version of this method (whereby listeners were only probed once at the end of each excerpt) to examine the occurrence and nature of listeners’ mind wandering experiences during music listening. The authors found that, compared to the experience of words, visual imagery was a prevalent aspect of participants’ mind wandering experiences. Another study using experience sampling to examine mind wandering during music listening in everyday life showed that visual imagery was occurring about 19% of the time, with up to 65% of music-related mind wandering occurring spontaneously (Taruffi, 2021). Such studies have been valuable in providing some indications of the incidence of music-evoked imagery and mind wandering including with respect to its intentionality. However, studies that use the probe-caught methodology to examine the rate of occurrence (Weinstein, 2018), the patterns of intentionality and the neural correlates (Polychroni et al., 2022) of visual imagery over the course of a single music listening episode are still lacking.

### 1.2. Neural correlates of music-evoked visual imagery

A growing body of work shows that while seemingly highly subjective, the occurrence and the contents of visual imagery can nevertheless be seen reflected in neural activity (Cooper et al., 2003; Dijkstra et al., 2017, 2019; Drever, 1955; Klimesch et al., 2007; Williamson et al., 1997; Xie et al., 2020). The study by Hashim et al. (2024) confirmed that music-evoked visual imagery is associated with suppressed alpha power [8-13 Hz], and in doing so, corroborated findings from the only other neuroscience study to examine music-related visual imagery (Fachner et al., 2019). However, in addition to attenuated alpha power in primarily posterior regions of the brain, research has also implicated other potential correlates of music-evoked visual imagery. For example, gamma power in the occipital brain area has been associated with spontaneous imagery conjured up during deep meditation (Luft et al., 2019). Similarly, in addition to theta having been shown to aid discrimination of the contents of visual imagination (Xie et al., 2020), theta and delta power in frontal, central, and parietal areas, have been associated with mind wandering (Polychroni et al., 2022). The latter finding may be particularly relevant given that visual imagery is sometimes considered a form of mind wandering (Taruffi, 2021; Taruffi et al., 2017).

Hashim et al.’s (2024) study showed that while both static and dynamic imagery led to alpha suppression during visual imagery formation as expected (Klimesch, 1999; Klimesch et al., 2007; Sousa et al., 2017; Xie et al., 2020), dynamic imagery was further associated with brain areas and oscillatory bands typically linked with motoric action and higher internal processing demands (De Lange et al., 2008; Menicucci et al., 2020; Sepúlveda et al., 2014; Zabielska-Mendyk et al., 2018). It is thus relevant to ask how alpha and other neural oscillations may also differ as a function of whether one is experiencing visual imagery spontaneously or due to deliberate action.

With regard to alpha, whether spontaneous visual imagery is reliably associated with alpha suppression remains an open question. On the one hand, while previous research suggests that spontaneous thought is prominent in the listening experience (Taruffi, 2021), the one previous paradigm to systematically examine the neural correlates of music-evoked visual imagery may have (by continually asking participants *how much* imagery was experienced) largely encouraged deliberate imagery (Hashim et al., 2024). On the other hand, since spontaneous imagery is more likely to be the result of automatic lower-level visual processing (in contrast to deliberate imagery which is more likely the result of top-down mechanisms^1^; for a review, see Pearson, 2019), one possibility is that posterior alpha suppression, which reflects activity in occipital areas would be even more prominent for spontaneous than for deliberate imagery.

Whether differences between spontaneous and deliberate imagery may be expected outside the alpha band is also unclear. Findings of higher theta power during unaware mind wandering states have been linked to similar findings during meditative and absorption states (Polychroni et al., 2022). Similarly, delta power increases found during unaware mind wandering have been interpreted as indexing the maintenance of internal trains of thought through the inhibition of external interference (Polychroni et al., 2022). Given that visual imagery may be considered a form of mind wandering, one prediction is that, as for unaware mind wandering, spontaneous visual imagery will be associated with enhancements of both delta and theta activity. However, since intentionality of visual imagery (spontaneous experiences vs. deliberate experiences) is only partially related to the metacognitive awareness of imagery (i.e., spontaneous imagery, due to the lesser cognitive control involved, may be expected to be associated with lesser awareness than deliberate imagery, but may still reach conscious awareness), expecting similar patterns of activity as in the study from Polychroni and colleagues may not be wholly justified.

Taken together, research has provided evidence that music-evoked visual imagery has reliable neural correlates. However, still unclear is if and how the neural correlates of spontaneous forms of visual imagery during music listening and in general, may differ from neural correlates of more deliberate imagery. Such insights would improve not just understanding of music-evoked visual imagery but also understanding of how the brain mediates mental actions that differ in their intentionality.

### 1.3. The present research

The current experiment sought to examine the incidence and neural correlates of spontaneously and deliberately occurring visual mental imagery in response to music. Using an ecologically valid probe-caught method (Polychroni et al., 2022; Taruffi et al., 2017; Weinstein, 2018), participants were regularly probed on both visual imagery occurrence and on the intentionality of any such imagery throughout extended music listening trials. Specifically, using dichotomous responses, they reported on whether they had or had not experienced any visual imagery immediately prior to receiving a probe (providing listeners with the option to report no visual imagery at all), and, if answered in the affirmative, whether the formation of this imagery had been spontaneous or deliberate. Musical stimuli differing in terms of relaxation potential (relaxing vs. non-relaxing) as well as familiarity (experimenter-selected vs. participant-selected) were used to allow generalisation of findings to the wide range of stimulus types that are typically experienced in everyday life (Jakubowski & Francini, 2022; Pereira et al., 2011). Using stimuli varying in familiarity and relaxation potential also allowed investigation of how these factors influence the relative incidence of spontaneous versus more deliberate forms of visual imagery. Finally, individual differences in musical training, general imagery ability and music preference were measured to allow examination of if and how these may influence imagery experiences.

In light of the existing research and gaps in research, we hypothesised the following:

1. All music-evoked visual imagery will be characterised by alpha power suppression found primarily in the posterior regions of the brain (Fachner et al., 2019; Hashim et al., 2024; Sousa et al., 2017; Xie et al., 2020).
2. Spontaneous imagery will be associated with alpha power suppression (in line with previous work that has linked alpha suppression to visual imagery, albeit not with spontaneous imagery specifically) and may even be to a greater degree than for deliberate visual imagery (this due to the latter being a more top-down process and, as such, a potentially less vivid visual experience-led phenomenon; Seli, Risko, Smilek, et al., 2016).
3. Spontaneous visual imagery may be associated to some extent with increases in both theta and delta power; this, because spontaneous imagery may be less likely to reach conscious awareness and may thus be more similar (than deliberate visual imagery) to unaware mind wandering (Polychroni et al., 2022).
4. Familiar music may be associated with a greater incidence of music-evoked visual imagery experience (e.g., due to its greater association with autobiographical memory), as may a higher overall general visual imagery ability of the listener and a greater preference for the genres of the musical stimuli in question.

## 2. METHODS

### 2.1. Participants

Data was collected from 30 participants (aged 18-49, 20 female, 9 male, 1 non-binary; M = 27.03, SD = 7.5) who were recruited using poster advertisements, word-of-mouth, and a student credit scheme. We deemed our sample size sufficient as research into the correlates of music-evoked visual imagery have shown expected effects in a single participant case study (Fachner et al., 2019).

An online survey was used to assess eligibility of participants for the current research. The survey asked participants to report (on a 1-7 Likert scale, from 1 = ‘Not at all’ to 7 = ‘Experience very strongly’) the extent to which they experienced the following phenomena in response to music: chills or goosebumps, visual imagery, felt emotions, and personal memories (where the selection of experiences were probed in order to help mask the intentions of the study). The primary inclusion criterion was that individuals report experiencing at least mild levels (providing a rating of 2 or upwards) of visual imagery in response to music. Those who reported never experiencing music-evoked visual imagery (i.e., provided a rating of 1) were considered ineligible and were thanked for their time.

This research was approved by the Research Ethics Committee of the Department of Psychology at Goldsmiths, University of London. All participants provided online and in-person written informed consent before taking part. Eligible participants were provided with monetary compensation of £10 per hour for their time.

### 2.2. Obtaining self-selected stimuli

In addition to screening participants, the initial online survey asked eligible participants to provide examples of musical pieces that they tended to listen to in order to relax as well as up to four examples of pieces that they would *not* choose to listen to when looking to relax. These questions were modelled after those used by Baltazar et al. (2020). Specifically, the first question asked: ‘*Imagine you are feeling anxious, stressed or nervous, but you need to calm down in order to be able to focus on your work. Whilst in this situation, you decide to listen to some music to help you relax. What would be some good examples of music pieces that would work for you in this kind of situation?*’. The second question asked: ‘*Please also think of music pieces you are familiar with and you like, but would not work well in this stressful situation. In other words, music that would not necessarily help you feel more relaxed.*’ Participants were encouraged to provide 2-4 music examples for each question and were asked to provide examples not containing lyrics or (where this was not possible) to provide acoustic alternatives to the lyrical versions.

### 2.3. Pilot survey for the selection of unfamiliar music

A pilot survey was conducted to obtain experimenter-selected musical excerpts for the main experiment (the full results of this pilot survey are provided in Supporting Information). We recruited a total of 17 participants (aged 18-42, female = 10, male = 7, M = 26.71, SD = 7.04), and the survey included excerpts from electronic, classical, and jazz genres. The tracks were selected from a range of sources, but mostly from Marti-Marca et al. (2020), allowing us to vary pieces on the basis of arousal as a basic index of the relaxation potential of a given track.

In total, twelve pieces were used in the pilot survey (see Supplementary Table 1 in Supporting Information for the full list): six that were considered high in relaxation potential and six low in relaxation potential with the inclusion of two options from each genre (EDM, classical, and jazz) within each group. Participants rated (using a 1-7 Likert scale) parameters similar to those used by Baltazar and Västfjäll (2020), namely: energy (how intense or active the music was), acousticness (how much the track consisted of acoustic instruments or electronic sounds), valence (the positivity or negativity of the music), danceability (suitability for dancing), and loudness (overall volume conveyed). Participants were also asked how relaxing they found the music, how much they liked it, and how familiar they were with it.

The final stimuli set used in the main experiment is indicated in Supplementary Table 1 in Supporting Information in further detail. To be labelled as having high relaxation potential, tracks needed to be rated on a 1-7 Likert scale as having low (ratings of 3 or below) energy, low danceability, low loudness, and high (ratings of 4 and above) relaxingness on average. In contrast, low relaxation potential tracks needed to be rated with high energy, high danceability, high loudness, and low relaxingness. Acousticness was used to confirm that an equal number of acoustic and electronic tracks were included in the two groups. Valence and liking ratings were collected to ensure that tracks were rated as being at least neutral or higher in terms of positivity and enjoyment. Familiarity ratings were obtained to ensure that tracks selected for the two groups were similarly unfamiliar (presented as ‘*How familiar are you with this excerpt?*’ and rated along a 4-point Likert scale from 1 = ‘As far as I know, I’ve never heard this excerpt before’ to 4 = ‘I know this excerpt, it is called/by: [*open-text response*]’). As observed in Supplementary Table 2 in Supporting Information, familiarity values were found to be generally low across all music tracks, confirming their unfamiliarity.

### 2.4. Materials and stimuli

#### Listening conditions

The listening task was comprised of four conditions: an unfamiliar (i.e., experimenter-selected) relaxing, an unfamiliar non-relaxing, a familiar (i.e., participant-selected) relaxing, and a familiar non-relaxing music condition. In the event that the familiar music stimuli provided by participants in the screening survey did not together extend to 20 mins in duration (the full length of each listening trial), excerpts were repeated, starting from the longest track to the shortest (in order to avoid too many repetitions of the participant’s music), until the full 20-min duration was reached.

#### Probe-caught measure

Visual imagery experience was assessed by presenting thought probes at pseudorandom intervals throughout the listening trials. Probes were presented every 40 s with a uniform ± 10-s jitter (i.e., all possible durations between 30 and 50 s across every listening condition). Phrasing and implementation of the thought probes followed previous mind wandering literature (Polychroni et al., 2022; Taruffi et al., 2017; Weinstein, 2018). The first probe prompted the participant to indicate if they were experiencing any visual imagery: ‘*Just before the probe, were there any images in your mind’s eye?*’, presented with the dichotomous response boxes: ‘*Yes’* and ‘*No’*. If the participants responded with ‘*Yes’*, they were asked a follow-up question: ‘*Were your thoughts deliberate and actively generated or spontaneous and free-flowing?*’, and then presented with the dichotomous response boxes: ‘*Deliberate’* and ‘*Spontaneous’*. Participants were given up to 4 s to answer each probe. These parameters were set to allow presentation of as many probes as possible (a total of up to 25 probes per listening condition and up to 100 probes over the four conditions), while allowing sufficient time in between probe presentations for the participant to return to the listening task. The limited time given to respond was also to discourage participants from overthinking their answers and to avoid interrupting their listening experience for too long each time.

#### Affect ratings

At the end of each listening condition, we collected affect ratings on a visual analogue 1-7 Likert scale (‘*Please rate the following indices to best reflect your current state*’) with questions similar to those used by Baltazar et al. (2019): valence (negative– positive), energy (drowsy–alert), and tension (relaxed–tense). An additional thirteen questions further inquiring into the content of visual imagery experienced as well as general impressions regarding the music heard during the listening trials were also presented at the end of each block (see Supplementary Table 3 in Supporting Information for a list of these additional questions). However, this data will be analysed and discussed elsewhere.

#### Individual differences

The 7-item Musical Training subscale of the Goldsmiths Musical Sophistication Index (Gold-MSI; Müllensiefen et al., 2014) was presented to gauge participants’ levels of prior musical training (e.g., ‘I have never been complimented for my talents as a musical performer’, ‘I am able to judge whether someone is a good singer or not’), using a 7-point Likert scale from 1 = ‘Completely Disagree’ to 7 = ‘Completely Agree’.

We further presented the 5-item Vision and 5-item Emotion subscales of the Plymouth Sensory Imagery Questionnaire (Psi-Q; Andrade et al., 2014), which assesses one’s general ability to experience mental imagery across multiple modalities (such as visual, tactile, emotion, olfactory, etc.; e.g., ‘Imagine the appearance of … a sunset’, ‘Imagine feeling … excited’). These items were rated on an 11-point Likert scale from 0 = ‘no image at all’ to 10 = ‘imagery as clear and vivid as real life’.

Finally, a selection of genres (classical, electronic dance music, jazz, and ambient) were presented to examine participants’ preferences for the musical genres presented within the experimenter-selected tracks (‘*Please indicate your basic preference for each of the following genres using the scale provided.*’). Participants rated their preference for each genre using a 7-point Likert scale from 1 = ‘Dislike Strongly to 7 = ‘Like Strongly’.

### 2.5. Procedure

The experiment was programmed and presented using the experiment-building software, OpenSesame (Version 3.3.8, Python 3.7.9; Mathôt et al., 2012). Participants sat in a dimly lit room and were presented with verbal as well as written instructions. The stimuli were played through headphones and participants were advised to adjust the volume to a comfortable level while listening to a sample listening track and to maintain that volume for the duration of the experiment. Participants completed – in randomised order – the four listening blocks with music differing in familiarity and relaxation potential, and with each block lasting approximately 25 mins. The full duration of the experiment was between 2 and 3 hours (including EEG preparation, verbal and written instructions, and potential questions). Prior to the start of the experiment, deliberate visual imagery was defined as *visual thoughts that are purposefully thought out*, whereas spontaneous visual imagery was defined as *appearing without any particular intention or control*.

For each of the four listening blocks, participants first stared at a fixation dot for 30 s, before being asked to provide ratings (using three affect ratings) on how they were feeling at that moment and then being presented with the 20-minute music block. Participants were informed that they should listen to the music presented, by immersing themselves with eyes closed, and that they would hear a brief clicking sound any time a probe question appeared on the screen. Participants were told they should open their eyes at that point, provide their answer, and then continue to listen with eyes closed. At the end of each block (indicated by a final click sound), participants were once more required to fixate for 30 s and to complete the affect questions, before answering additional questions about their visual imagery experience. To finish, participants filled in the Musical Training subscale of the Gold-MSI, the Vision and Emotion subscales of the Psi-Q, and the short musical genre preference survey.

### 2.6. EEG recording and analyses

EEG was recorded using the mobile Waveguard 32-channel cap from ANT Neuro, The Netherlands. The 32 channels used were: Fp1, Fpz, Fp2, AFz, F7, F3, Fz, F4, F8, FC5, FC1, FC2, FC6, T7, C3, Cz, C4, T8, M1, CP5, CP1, CP2, CP6, M2, P7, P3, Pz, P4, P8, POz, O1, Oz, O2. The cap was placed in accordance with ANT Neuro’s 5% electrode placement system, a derivative of the 10-20 international system. Signals were amplified using the eego EEG recording amplifier, with the reference set to electrode CPz. Impedance levels were kept low and never above 20 kΩ. The EEG recording was set to a sampling rate of 500 Hz for all participants.

The EEG data was imported into MATLAB, pre-processed using EEGLab (Delorme & Makeig, 2004), and further analysed using functions from the FieldTrip toolbox (Oostenveld et al., 2011). Data segmentation, pre-processing and analysis choices were broadly in line with previous studies (Compton et al., 2019; Liu et al., 2021; Polychroni et al., 2022). The continuous data was first segmented into 34.7-s epochs: –12 s before the probe onset to 22.7 seconds after probe onset. The time window of interest was from –10 to 0 s relative to probe onset. However, the longer epoch was taken to allow a 700-ms baseline window within the interstimulus interval (after probe onset and before the start of the next probe), as well as to avoid edge artefacts.

The segmented data was resampled to a rate of 200 Hz. Further, a low-pass band filter was set at a frequency of 50 Hz and a notch filter at a frequency of 48-52 Hz. Bad or faulty electrodes were visually searched for by observing the raw data, but as none were found, no electrodes were removed or interpolated in this step. Artefacts were then visually assessed by scrutinising the time courses of each trial’s raw data and replacing bad segments with NaN’s (average seconds removed = 1.31, max = 6.32, min = 0.08). Next, independent component analysis (ICA) was used to identify and remove artefacts caused by eye movements, eye blinks, heartbeats, and further channel noise (average number of components removed = 4.7). Finally, trial ✖ channel spectra were visually assessed to reject whole trials that still exhibited high levels of variance even after prior artefact removal efforts (average number of trials excluded = 0.93). All the data was then re-referenced to the average of all the electrodes.

Finally, in preparation for statistical analyses, epoched data was subdivided into four groups corresponding to all possible visual imagery states. These groups were *no visual imagery*, *all visual imagery*, *deliberate* visual imagery only, and *spontaneous* visual imagery only.

### 2.7. Time-frequency decomposition

A time-frequency decomposition of epochs was computed using a Hanning taper moving with 50 ms steps along the 34.7-s epoch for frequencies from 1 to 50 Hz. This decomposition was carried out using a frequency-dependent window length, with 7 cycles per time window. Spectral power for every trial was averaged then normalised by dividing it by the baseline level (i.e., the gain model; Grandchamp & Delorme, 2011).

A 700-ms baseline was selected as 15 to 15.7 s post probe onset, as it was late enough in the inter-probe interval to be sure that participants had closed their eyes again after responding to a probe (participants had up to 8 seconds to respond to the two questions) but early enough to avoid that substantial visual imagery had been formed again. Importantly, comparing this choice of baseline period against one that was earlier (10 s after probe onset) and another that was later (20 s after probe onset) showed that the nature of effects reported below (see section *4.1 Oscillatory characteristics of music-evoked visual imagery*) did not differ across the three baseline windows.

## 3. STATISTICAL ANALYSES

### 3.1. Spectral analysis: differences between visual imagery manifestations

Differences in the spectral power between the different visual imagery states (visual imagery vs. no visual imagery, spontaneous vs. deliberate imagery, spontaneous vs. no visual imagery, and deliberate vs. no visual imagery) were analysed using non-parametric cluster-based paired permutation tests (Maris & Oostenveld, 2007). These were run for each of the oscillatory frequency bands of interest (delta [2-3 Hz], theta [4-7 Hz], alpha [8-13 Hz] and gamma [30-45 Hz] for completion) using functions from the MATLAB-based Fieldtrip toolbox (Oostenveld et al., 2011).

These analyses were first carried out by calculating the observed test statistic by running paired samples t-tests between the specified contrasts at each sample of the 3-dimensional data (channel vs. frequency vs. time). Those samples for which the t-statistic fell beneath the **α** < 0.05 threshold were selected and clustered in connected sets on the basis of temporal vs. spatial vs. spectral adjacency. The t-statistics within every cluster were then summed to calculate the cluster-level statistics. The maximum of these statistics served as the test statistics with which differences between visual imagery and manifestation states were determined.

Subsequent calculation of the cluster significance probability was done using the Monte Carlo method. The previous steps were repeated once more but calculated for 500 permutations of randomly partitioned data in two subsets. These values were then compared to the observed test statistic. Significance levels pertaining to effects in the alpha band were set at a *one*-tailed **α** < 0.05 (and a minimum of two neighbouring channels), due to consistent findings of posterior alpha suppression in response to moments of general and music-related visual imagery (Dijkstra et al., 2019; Fachner et al., 2019; Hashim et al., 2024; Sousa et al., 2017; Xie et al., 2020). All other cluster significance levels were set at a *two*-tailed **α** < 0.05, with a minimum of two neighbouring channels considered as a cluster.

Participants varied in their visual imagery and intentionality probes, which resulted in contrasting number of trials for each intentionality state within each comparison. Discrepancy scores (%) were calculated between states within contrasts for each participant, and those with a 100% discrepancy score (i.e., whereby a participant did not report experiencing one or the other state) were excluded. Four main contrasts were implemented with mostly variable trial numbers (M ± SD) and sample sizes [N]: (1) visual imagery (55.4 ± 30.4) vs. no visual imagery (21.4 ± 18.3) [25]; (2) spontaneous imagery (36.9 ± 23.4) vs. no visual imagery (21.4 ± 18.3) [25]; (3) deliberate imagery (18.5 ± 12.9) vs. no visual imagery (21.4 ± 18.3) [25]; (4) spontaneous imagery (43.8 ± 22.3) vs. deliberate imagery (20.8 ± 13.2) [28]. Where relative discrepancy scores appeared highly in favour of one state within a comparison, we performed a set of control analyses to systematically minimise discrepancies as much as possible while retaining as many participants as we could (see *Control subset analyses: cluster-based permutation analysis* in Supporting Information). To foreshadow the results, these control analyses showed that the main claim of the manuscript (namely that spontaneous imagery results in reliable alpha suppression) was not an artifact of differences in trial numbers.

### 3.2. Behavioural analyses

The statistical analyses of the probe data and affect ratings were carried out using R (Version 4.2.3; R Core Team, 2018). All subsequently defined linear mixed-effect models (LMMs) were computed using the *lme4* (Bates et al., 2015) and *lmerTest* (Kuznetsova et al., 2017) R packages.

#### 3.2.1. How affect changes as function of music’s qualities

LMMs were run to check that exposure to music with different qualities had effects on the listeners’ affect as might be expected. Specifically, we defined a separate model with each affect rating (energy, tension, and valence) as a dependent variable, and with categorical variables Time Point (Pre vs. Post), Familiarity type (Familiar vs. Unfamiliar), and Relaxation Potential type (Relaxing vs. Non-Relaxing) as a fixed effects, as well as participant ID as a random effect.

#### 3.2.2. Influence of music qualities and individual differences (musical training, imagery ability, and preference for genres of heard music) on visual imagery incidence rate

Potential differences in the incidence rates (%) of visual imagery occurrence (*yes* vs. *no*) and intentionality (*spontaneous* vs. *deliberate*) as a function of music qualities and individual differences were estimated with LMMs. In a first model, the dependent variable was visual imagery occurrence (*yes* vs. *no*) rates for each participant for each block while Familiarity (familiar vs. unfamiliar), and Relaxation Potential (relaxing vs. non-relaxing) were fixed effects with an interaction term specified between them. The second model further included Musical Training, the two Psi-Q dimensions (Vision and Emotion) and mean preference ratings across all used genres (EDM, Classical, Jazz, and Ambient) as fixed effects. Participant ID was included as a random effect in all models. The model assessing visual imagery intentionality occurrence rate was similarly designed, however using rate of occurrence of *spontaneous visual imagery* (as opposed to deliberate) as the dependent variable.

## 4. RESULTS

Participants reported experiencing visual imagery in response to 72.59% (SD = 22.17) of probes across all listening conditions. Thus, visual imagery was reported more often than *not* experiencing visual imagery (M = 21.15%, SD = 40.85; note that 6.26%, SD = 24.23, pertained to missed probes). Participants also reported experiencing spontaneous visual imagery (M = 70.94%, SD = 45.41) more often than deliberate visual imagery (M = 28.84%, SD = 45.31).

### 4.1. Oscillatory characteristics of music-evoked visual imagery

#### 4.1.1. Alpha band

As can be observed in Figure 1, the cluster-based permutation tests revealed a significant suppression of alpha power in the comparison between **visual imagery and no visual imagery** probes occurring from –4.8 to –3.5 s prior to probe onset, *p* = .030. The effect was topographically dispersed, located around midline fronto-central channels during the initial stages, before moving to right parieto-occipital and midline occipital channels by the end of the time window (see Figure 1b and 1d).

**Figure 1.**
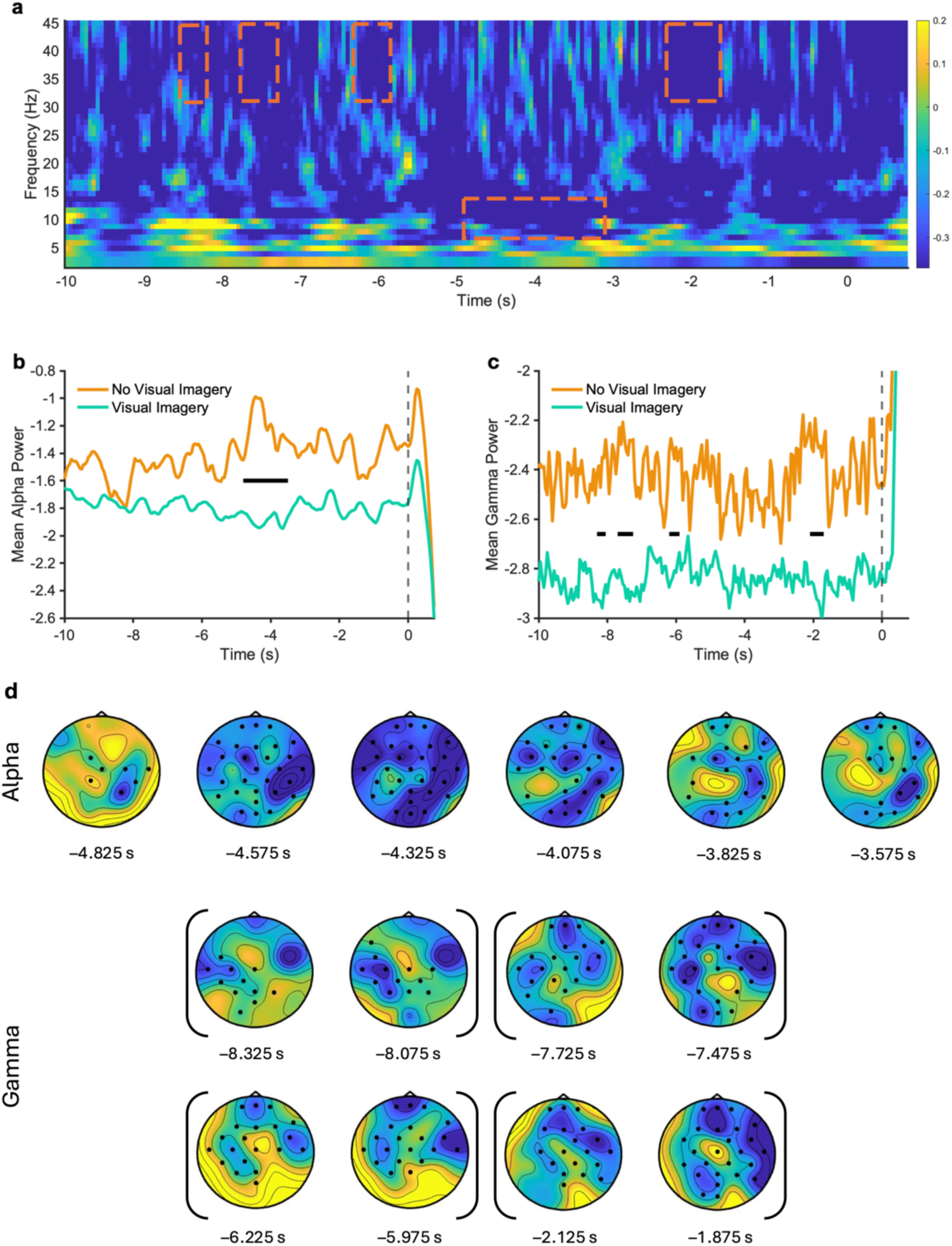
Oscillatory differences between states (visual imagery and no visual imagery) as a function of time (s) relative to probe onset (0 s; indicated in the line plots by a dotted line along the y-axis) for alpha [8-13 Hz] and gamma [30-45 Hz] bands. (a) Time-frequency spectrogram averaged across channel sites of the power difference between visual imagery and no visual imagery probes. Clusters reflecting significant state differences are denoted by broken orange rectangles. (b) Alpha spectral power averaged over the channel sites of the cluster. The significant cluster is denoted by the black bar along the x-axis. (c) Gamma spectral power averaged over the channel sites of the clusters. The significant clusters are denoted by the black bars along the x-axis. (d) Topography of the clusters within each frequency band at 250 ms intervals (black markers denote electrodes that were present in the cluster). Rounded brackets surrounding sets of topoplots represent individual cluster groups.

Analyses further revealed alpha suppression between **spontaneous and no visual imagery** probes in three clusters: (1) –1.1 to 0 s time range, *p* = .046; (2) –3.1 to –1.5 s time range, *p* = .048; and (3) –5.25 to –3.4 s time range, *p* = .012 (see Figure 2a). These clusters were also topographically dispersed across sites, but greatest alpha suppression across all three clusters tended to be located in midline and left parieto-occipital channels (Figure 2c and 2g).

**Figure 2.**
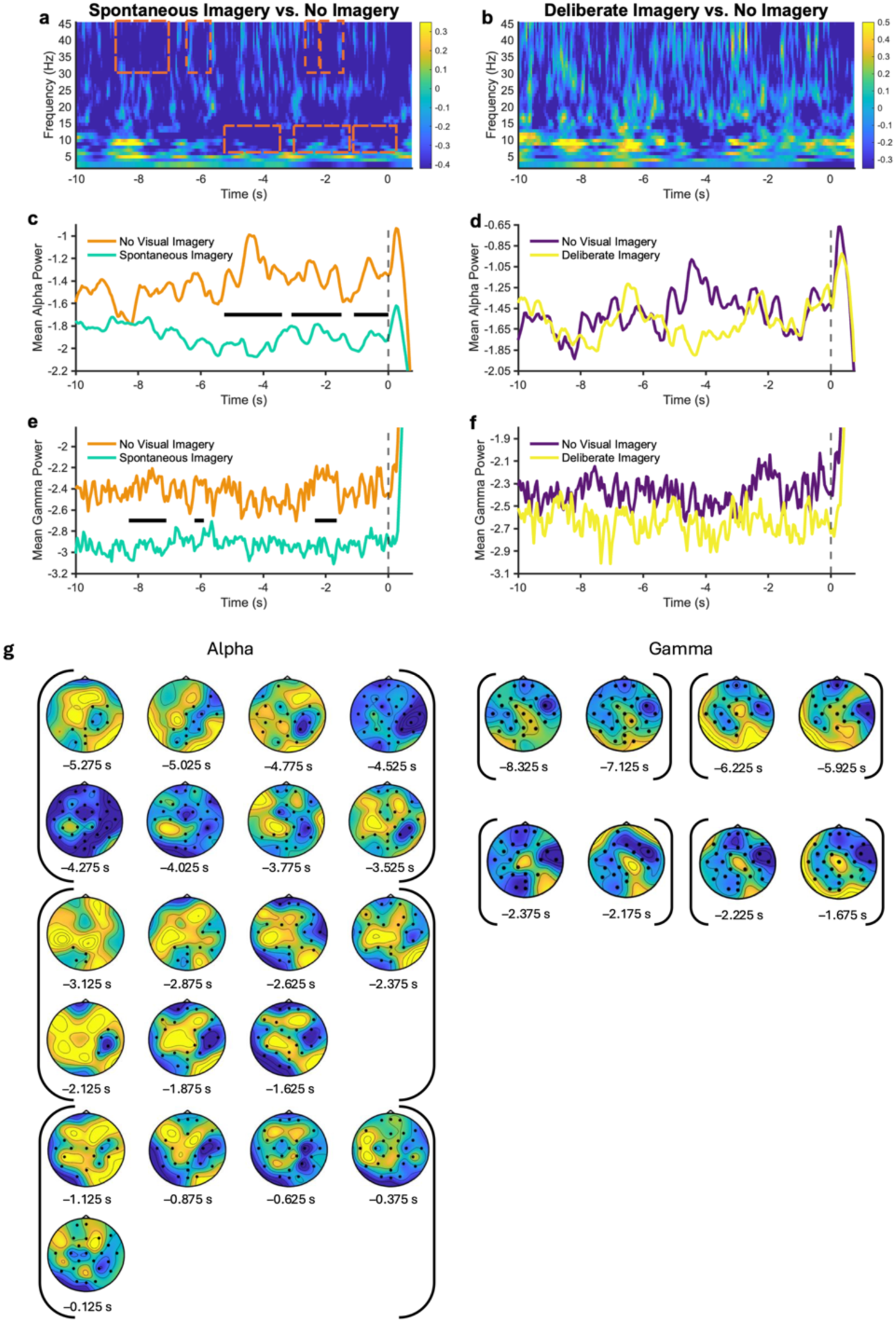
Oscillatory differences between states (spontaneous/deliberate imagery and no visual imagery) as a function of time (s) relative to probe onset (0 s; indicated in the line plots by a dotted line along the y-axis) for alpha [8-13 Hz] and gamma [30-45 Hz] bands. (a) Time-frequency spectrogram averaged across channel sites of the power difference between spontaneous imagery and no visual imagery probes. Clusters reflecting significant state differences are denoted by broken orange rectangles. (b) Time-frequency spectrogram averaged across channel sites of the power difference between deliberate imagery and no visual imagery probes. No significant differences were found. (c/e) Alpha/Gamma spectral power averaged over the channel sites of the clusters reflecting state differences between spontaneous and no visual imagery states. The significant clusters are denoted by the black bars along the x-axis. (d/f) Alpha/Gamma spectral power averaged over all channels between deliberate and no visual imagery states. (g) Topography of the clusters reflecting state differences between spontaneous and no visual imagery states within each frequency band. Alpha clusters were segmented at 250 ms intervals (black markers denote electrodes that were present in the cluster), whereas gamma clusters were plotted in their original time windows due to their short-lived nature. Rounded brackets surrounding sets of topoplots represent individual cluster groups.

In contrast, the differences between **deliberate and no visual imagery** states (*p* = .367) and between **spontaneous and deliberate imagery** states in the alpha band were not significant (*p* = .078).

#### 4.1.2. Delta, theta, and gamma bands

Analyses revealed no differences between any of the visual imagery states in terms of delta and theta power. Specifically, no significant differences were found between visual imagery and no visual imagery probes, between no visual imagery and spontaneous and deliberate imagery probes, as well as between spontaneous and deliberate imagery probes.

However, **visual imagery (compared to no visual imagery)** was associated with reduced gamma power as visible in four temporally brief clusters (see Figure 1a and 1c; cluster 1: –2.1 to –1.7 s, *p* = .010; cluster 2: –6.2 to –5.9 s, *p* = .034; cluster 3: –7.7 to –7.25 s, *p* = .010; and cluster 4: –8.3 to –8.05 s, *p* = .046). Gamma suppression across all clusters was mainly observed in midline and bilateral frontal, central and fronto-central regions (although it transiently occupied midline occipital channels in clusters 1 and 3; see Figure 2d).

While there were no differences found between **deliberate and spontaneous** states (*p* = .054) or between **deliberate and no visual imagery** states (*p* = .174; for the latter contrast, see Figure 2b and 2f) in terms of gamma power, a similar pattern was obtained when comparing **spontaneous imagery and no visual imagery** states. Specifically, there was less gamma during spontaneous visual imagery found in four clusters (cluster 1: –2.2 to –1.65 s, *p* = .010; cluster 2: –2.35 to –2.15 s. *p* = .038; cluster 3: –6.2 to –5.9 s, *p* = .020; and cluster 4: –8.3 to – 7.1 s, *p* = .004), which were often observed in in midline frontal, central and fronto-central channels, while briefly occupying left parieto-occipital channels (see Figure 2g).

### 4.2. Checking the efficacy of affective state manipulation using the music stimuli

Musical stimuli differing in terms of relaxation potential (relaxing vs. non-relaxing) as well as familiarity (experimenter-selected vs. participant-selected) were used in order to allow generalisation of findings to the wide range of stimulus types that are typically experienced in everyday life (Jakubowski & Francini, 2022; Pereira et al., 2011). LMMs were run to check that exposure to music with different qualities had effects on the listeners as might be expected. See Figure 3 for visualisations of relationships separated by each listening block. For **Energy**, results revealed a significant main effect of time point, *F*(1, 203) = 6.39, *p* = .012, and a significant interaction between time point and relaxation potential, *F*(1, 203) = 17.08, *p* < .001. Following up this interaction effect revealed that energy ratings in response to the relaxing stimuli were significantly lower post listening (M = 3.28, SD = 1.39) than pre listening (M = 2.38, SD = 1.50), ß = –0.90, SE = 0.18, *t* = –5.05, *p* < .001, and that there was no difference in energy ratings between time points for the non-relaxing tracks, ß = 0.22, SE = 0.20, *t* = 1.08, *p* = .283. There were no other main effects or interactions, including those pertaining to music differing in familiarity.

**Figure 3.**
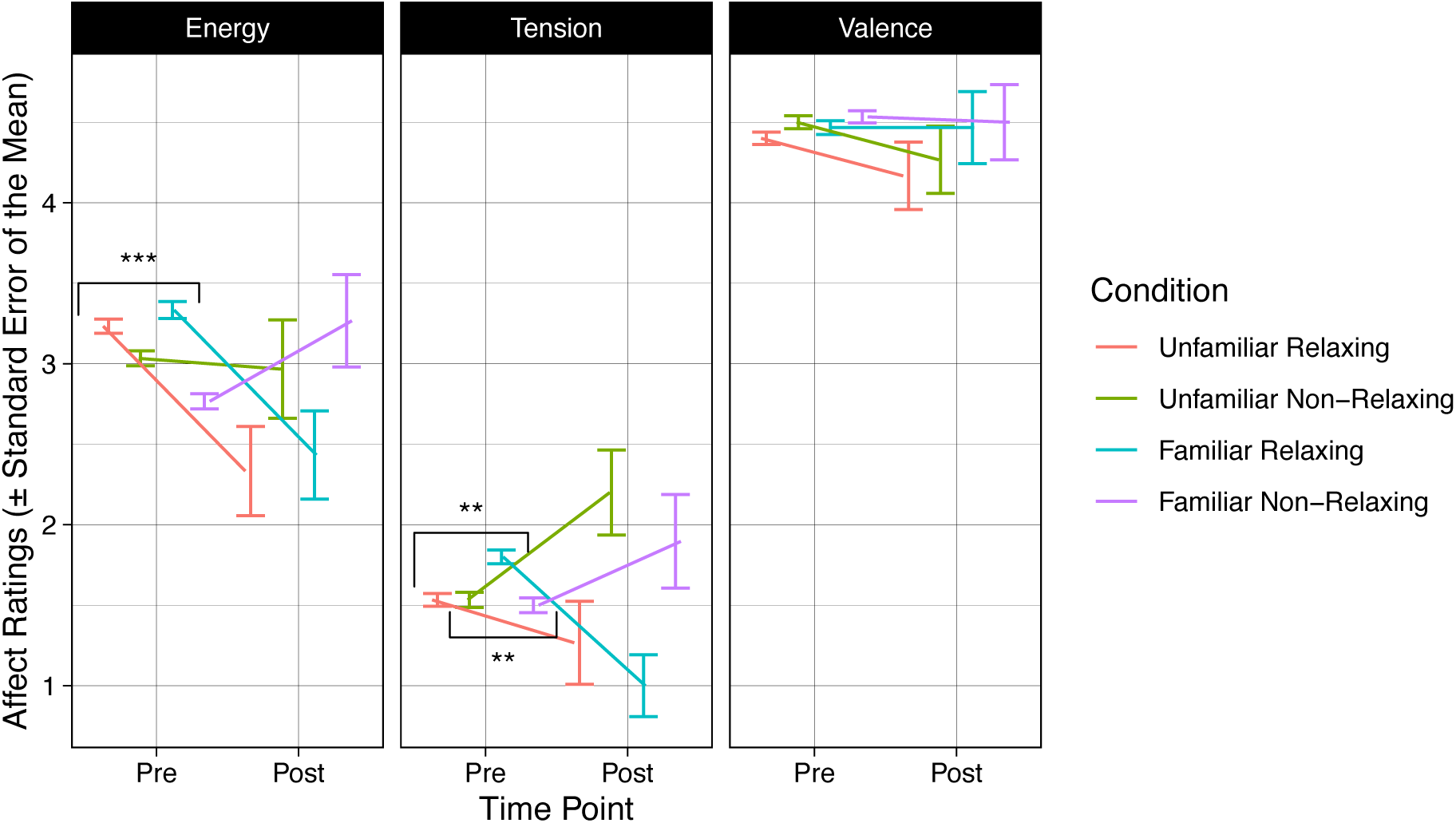
Mean values (and ± standard error of the mean) of affect ratings (Energy, Tension, and Valence) provided before (Pre) and after (Post) taking part in each listening condition (Unfamiliar Relaxing, Unfamiliar Non-Relaxing, Familiar Relaxing, and Familiar Non-Relaxing). Brackets indicate which music quality type (Familiarity vs. Relaxation Potential) yielded significant main effects between time points in the omnibus model. *** p < .001. ** p < .01.

For **Tension**, there was a significant main effect of relaxation potential, *F*(1, 202.2) = 8.29, *p* = .004, and a significant interaction between time point and relaxation potential, *F*(1, 202.2) = 16.26, *p* < .001. A follow up model showed tension ratings in response to the relaxing tracks to significantly decrease from the pre (M = 1.67, SD = 1.19) to post (M = 1.13, SD = 1.24) time points, ß = –0.53, SE = 0.17, *t* = –3.10, *p* = .003. In contrast, tension ratings increased from pre (M = 1.52, SD = 1.33) to post (M = 2.05, SD = 1.50) in response to the non-relaxing tracks, ß = 0.52, SE = 0.19, *t* = 2.68, *p* = .009. No other main effects or interactions were found, including those pertaining to music differing in familiarity. Regarding **Valence** ratings, we did not observe any significant main effects or interaction terms with regards to the effects of exposure to music differing in relaxation potential and familiarity on affect responses between time points.

Taken together, results confirmed that the music participants provided as having high or low relaxation potential modulated their affective states accordingly.

### 4.3. Exploring influences on the incidence of music-evoked visual imagery experience

Having confirmed the effectiveness of our stimuli manipulation in terms of arousingness/relaxation potential, a final set of analyses examined the influence of the different music conditions on the incidence of music-evoked visual imagery. LMMs predicting the incidence rate of visual imagery showed there to be a significant influence of Familiarity, ß = 0.07. SE = 0.03, *t* = 2.66, *p* = .009, and a significant influence of Relaxation Potential, ß = 0.06. SE = 0.03, *t* = 2.12, *p* = .036, whereby familiar tracks and relaxing tracks were both associated with higher visual imagery rate (see Figure 4). The interaction was not significant, ß = 0.05. SE = 0.04, *t* = 1.45, *p* = .150. The analysis looking at spontaneous and deliberate visual imagery alone did not result in any significant main effects or interactions, suggesting that neither familiarity nor relaxation potential influenced the rate at which spontaneous imagery is experienced relative to deliberate imagery.

**Figure 4.**
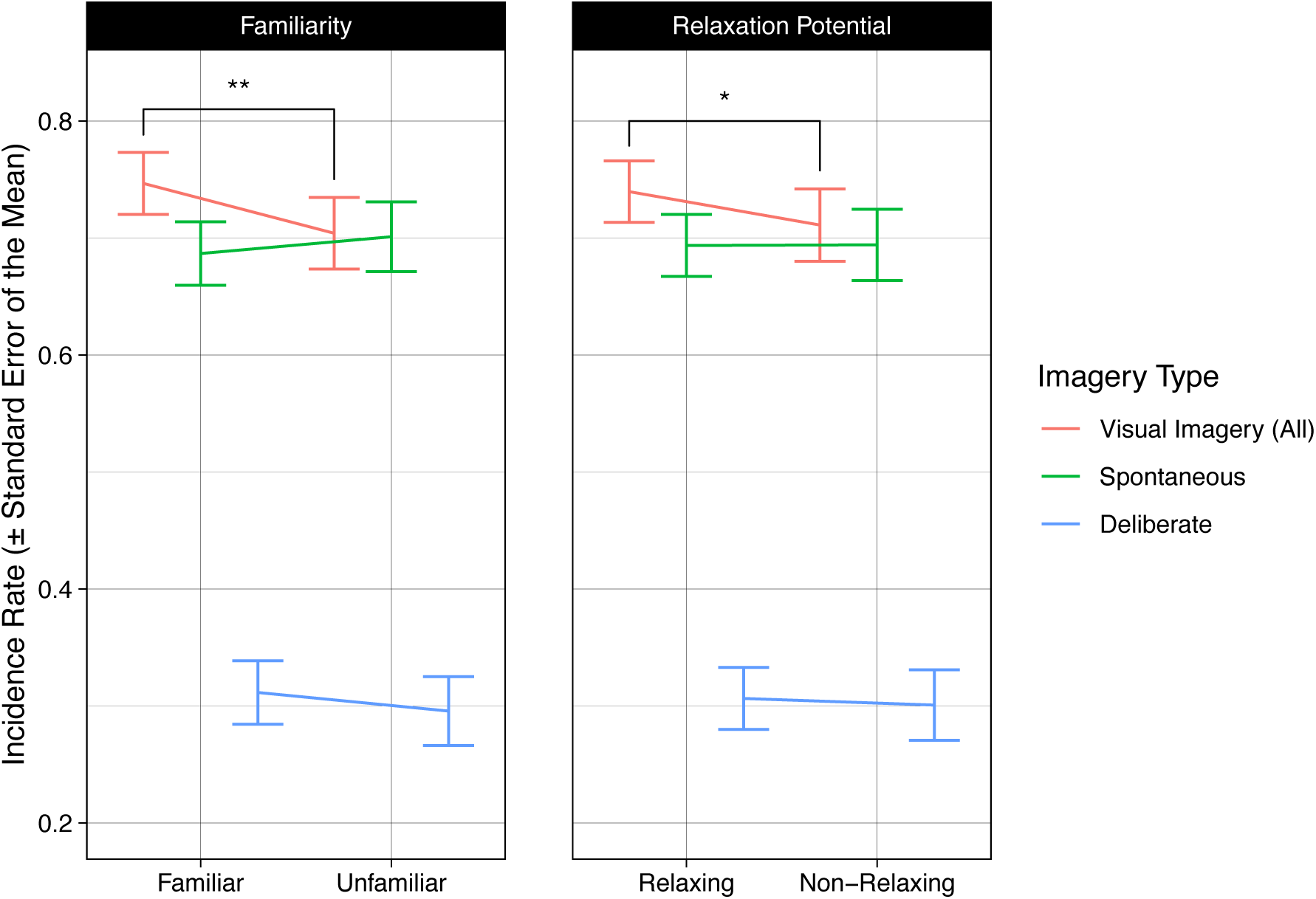
Mean incidence rate values (and ± standard error of the mean) of imagery ratings (Visual Imagery (All), Spontaneous Imagery, Deliberate Imagery) provided in response to each music quality type (Familiarity and Relaxation Potential). Visual Imagery (All) values represent all possible imagery episodes, and the spontaneous and deliberate imagery values represent the rate of imagery for that category. Brackets indicate which music quality type (Familiarity vs. Relaxation Potential) yielded significant main effects between time points in the omnibus model. ** p < .01. * p < .05.

Finally, analyses predicting visual imagery and spontaneous visual imagery ratings with regard to their relationships with individual difference measures (musical training, imagery ability, and musical preferences) did not result in any significant main effects or interactions. See Supplementary Figures 4 and 5 for visualisations of these relationships.

## 5. DISCUSSION

The current research used a probe-caught method along with EEG recording to investigate the oscillatory characteristics of spontaneous and deliberate music-evoked visual imagery. The main objectives were (1) to assess whether previous findings of the neurophysiological patterns of music-evoked visual imagery could be replicated, and (2) to examine the oscillatory signatures of spontaneous as compared with deliberate visual imagery, since previous paradigms may have predisposed participants to carry out the latter. Based on past literature (Cooper et al., 2003; Dijkstra et al., 2019; Fachner et al., 2019; Hashim et al., 2024; Sousa et al., 2017; Xie et al., 2020), we predicted that music-evoked visual imagery would be associated with alpha power suppression, especially in posterior regions of the brain, which we further predicted may be induced to a similar, if not a greater extent, in spontaneous imagery than in deliberate imagery. It was further speculated that, as for unaware mind wandering (Polychroni et al., 2022), spontaneous imagery might be associated with enhanced theta and delta power, due to the greater likelihood for spontaneous imagery (compared to deliberate imagery) to lie below conscious awareness. Finally, we predicted that rates of visual imagery would be greater during familiar music (due to a greater potential for evoking autobiographical memories) and that rates of visual imagery may be influenced by individual differences.

### 5.1. Neural signatures of music-evoked visual imagery

In line with several previous studies, including the two music-related investigations of visual imagery available (Fachner et al., 2019; Hashim et al., 2024), we were able to provide evidence for alpha suppression in response to music-evoked visual imagery, and mainly in occipital and parieto-occipital regions responsible for visual processing (Dijkstra et al., 2017, 2019; Xie et al., 2020). Further, spontaneous imagery showed a reliable and strong alpha suppression that was not seen for deliberate imagery.

This activation of visual cortices for spontaneous but not deliberate imagery in the current study is in line with the postulation that deliberate imagery, due to its top-down nature, may be reported even when imagery is less pictorial and less vivid, and thus less able to invoke activity in low-level visual regions. However, here it is important to emphasise that the difference in degree of posterior alpha suppression when comparing spontaneous and deliberate conditions did not reach significance. This suggests that any differences between the neural patterns of spontaneous and deliberate imagery were not that great, and a relevant question is the extent to which the current pattern of findings is due to the particular listening conditions used here. Indeed, some interesting questions that present themselves are whether deliberate imagery’s vividness and, accordingly, capacity to induce visual cortex activation may be related to the amount of time spent listening to a given music track, the nature of the listening context, and situation and individual differences in the listener, none of which have been carefully manipulated in work to date.

The results did not reveal any modulation of activity in either theta or delta activity as a function of visual imagery intentionality. Literature on mind wandering had reported elevated theta and delta during unaware types of mind wandering (Polychroni et al., 2022) and it seemed reasonable to suggest that spontaneous visual imagery cognition would be more unaware than deliberate. Indeed, given proposals of visual imagery and mind wandering potentially falling under the same umbrella of spontaneous cognition (see Taruffi & Küssner, 2019), and given research showing significant intertwinement between both experiences (Christian et al., 2013; Deil et al., 2023; Taruffi, 2021; Taruffi et al., 2017), we had speculated that spontaneous visual imagery might be associated with enhanced theta and delta activity in frontal areas (Polychroni et al., 2022). However, our results did not support such a hypothesis and as previously noted, it is thus possible that linking intentionality with meta-awareness and visual imagery with mind wandering may be too crude as far as associations go.

Lastly, we found visual imagery to be associated with reductions in gamma power primarily in frontal and central areas. This was not an effect we hypothesised, and it is worthy of note that it did not survive our control analyses, which aimed to equalise the number of trials between conditions. However, frontal gamma has been associated with sensory and cognitive processing across a range of contexts and domains (e.g., Pérez-Acosta et al., 2024), and perhaps most pertinently in this case, gamma power in fronto-central areas has also been associated with feelings of stress (see Vanhollebeke et al., 2022, for a review). Since it has been suggested that music’s ability to reduce stress may be at least partly due to its ability to encourage processes related to autobiographical memory and mental imagery (Panteleeva et al., 2018), this finding could be cautiously interpreted as evidencing an association between visual imagery and reduced correlates of stress. However, given that the current study did not manipulate or measure stress levels, it is clear that further work would be needed to examine such possible relationships.

### 5.2. Incidence of and factors influencing spontaneous and deliberate visual imagery

Visual imagery experiences were shown to be prevalent in the current study with over 70% of probed moments being associated with imagery (Dahl et al., 2022; Hashim et al., 2023; Küssner & Eerola, 2019; Vuoskoski & Eerola, 2015). Our study also provided a window into the rates of different types of visual imagery intentionality (Taruffi & Küssner, 2019), showing a higher prevalence of spontaneous visual imagery in comparison to deliberate visual imagery. Critically, this finding coincides with recent evidence that mind wandering is more often spontaneous than not and very often occurs in the visual domain (Taruffi, 2021).

Past literature that has reported high incidence rates of music-evoked visual imagery has relied on traditional rating methods, which may have encouraged deliberately formed visual imagery. Experience sampling techniques, in contrast, offer participants the possibility of identifying instances throughout the listening experience in which visual imagery had not occurred. Our results show that the probe-caught methodology, which is often used in mind wandering research (Polychroni et al., 2022; Taruffi et al., 2017), is useful in tracking nuances in the experiences of visual imagery. In a study by Taruffi et al. (2017) using a probe-caught paradigm, visual imagery was found to be a prevalent aspect of participants’ experiences of music (as, for example, compared to the prevalence of the experience of words – understanding the semantic concept): a result we replicate here. Given the simplicity of the probe-caught method and the effectiveness with which it can measure changes in attentional states (Weinstein, 2018), there is a lot to recommend it in exploring the frequency, behavioural, and neural correlates of different types of visual imagery during music listening.

An additional aim of the current study was to relate visual imagery incidence rates with individual differences such as musical training, general abilities to emote and experience visual imagery, and preferences for the music genres used as stimuli. Previous studies suggest a relationship between visual imagery rates and trait visual imagery to various degrees (e.g., Hashim et al., 2020, 2023; Küssner & Eerola, 2019). This study did not show such an influence of these individual difference measures on (spontaneous and deliberate) visual imagery incidence rates. However, one point to consider is that the approach for examining general visual imagery ability used here differs from those used in previous examinations (i.e., the VVIQ; e.g., Küssner & Eerola, 2019), with one possibility being that the scale used here is a less sensitive measure of trait imagery abilities. Similarly, while the absence of a relationship between imagery incidence and emotion imagination would seem to go against the literature suggesting a close intertwinement of visual imagery and emotion induction (Day & Thompson, 2019; Juslin, 2013; Juslin & Västfjäll, 2008; Vroegh, 2019), it remains possible that while imagery may contribute to emotion felt during music listening, one’s general ability to imagine emotions does not necessarily translate into experiencing higher amounts of visual imagery during music listening.

### 5.3. Implications of the research

In addressing research questions surrounding the neural signatures of visual imagery, one important consideration is the demand characteristics that inevitably arise in designs that require participants to explicitly report on their experience. By studying intentionality with a probe-caught method, we were able to investigate for the first time the formation of music-evoked visual imagery in a more ecologically valid listening arrangement, thus getting closer to its natural occurrence in everyday life. We argue that future work on music-evoked visual imagery, which exploit this method, has the potential to generate research insights with higher external validity than has been attained thus far.

Here, it is also interesting to note that while visual imagery may be regarded as a form of mind wandering (Taruffi et al., 2017; Taruffi & Küssner, 2019), mind wandering has been associated with alpha enhancement (Polychroni et al., 2022) rather than the alpha suppression reported here and in other studies (Cooper et al., 2003; Fachner et al., 2019; Hashim et al., 2024; Xie et al., 2020). Previous literature has proposed that alpha enhancement during mind wandering reflects diminished perceptual processing and a detachment from the external world (Polychroni et al., 2022), ideas in line with the concept of cognitive ‘idling’ and reduced attentional processing (Cooper et al., 2003; Foxe & Snyder, 2011; Palva & Palva, 2007). This largely frontal activity is different from the emphasis on alpha as tracking more or less visual processing when found in the occipital lobe. Nevertheless, future work is necessary to better understand both the extent of the overlap and distinctions between visual imagery and mind wandering, and the degree to which observed alpha power patterns are reflective of the different processes.

Finally, previous research has suggested the potential for visual imagery in emotion regulation (Küssner & Eerola, 2019) and as an important tool in therapeutic settings (Karagozoglu et al., 2013). Interestingly, in a meta-analysis by Panteleeva et al. (2018), spontaneous autobiographical memories and mental imagery were suggested to play as critical a role in music-related stress and anxiety reduction as the acoustic features of the musical stimuli in question. Our finding of reduced frontal gamma power during visual imagery, which has previously been proposed as a marker of stress reduction (Minguillon et al., 2016), thus raises the interesting possibility of a link between music-evoked visual imagery and stress and anxiety reduction. However, future work is needed to explore this further.

## 6. CONCLUSION

The findings of this research corroborate past literature linking posterior alpha suppression with music-evoked visual imagery, while expanding it to show that spontaneously generated visual imagery during music listening has a high incidence rate and is reliably associated with alpha power suppression. Further work is needed to determine how different music listening contexts may influence the incidence and neural correlates of both spontaneous and deliberate imagery, with the latter needing particular attention given the current study’s failure to show any differences between it and the absence of imagery. To this end, we suggest that future work employ probe sampling methodologies, which allow a range of imagery experiences to be compared and contrasted, and which can be easily adapted to simulate real-life listening conditions.

## 7. ACKNOWLEDGMENTS

We would like to thank Dr David Baker for facilitating communications between the authors and MassiveMusic. We would further like to thank MassiveMusic for providing the Ambient track used in the current study (see Supporting Information) and for funding this research.

## 8. DECLARATIONS

### Data availability statement

data and analysis scripts are available on request.

## Ethics approval statement

this research was approved by the Research Ethics Committee of the Department of Psychology at Goldsmiths, University of London.

## Author contributions (CRediT)

SH: Conceptualization, Data curation, Formal analysis, Investigation, Methodology, Software, Visualization, Writing – original draft, Writing – review & editing.

DO: Conceptualization, Formal analysis, Funding acquisition, Methodology, Project administration, Software, Supervision, Writing – review & editing.

## Footnotes

1 Specifically, where in spontaneous imagery, it may be expected that images occur automatically, freely and largely unbidden, reports of deliberate forms of imagery may include situations in which any imagery that is successfully bidden is at least partly conceptual (i.e., less pictorial and potentially less vivid), and thus not as associated with visual cortical areas as spontaneous imagery is (Gilbert & Li, 2013; Mechelli et al., 2004).

