## Supporting Information for "SPONTANEOUS VISUAL IMAGERY DURING EXTENDED MUSIC LISTENING IS ASSOCIATED WITH RELIABLE ALPHA SUPPRESSION"

^a^ Corresponding author:

Sarah Hashim

Department of Psychology

**Pilot survey for the selection of unfamiliar music of high and low relaxation potential: *Continued***

We ran paired samples t-tests to examine the differences between the tracks included in the pilot study along a set of acoustic qualities (energy, acousticness, valence, danceability, and loudness) as well as a couple of aesthetic qualities (relaxingness and liking) and familiarity, for the purpose of selecting a final set of tracks to be presented as part of the main study’s unfamiliar relaxing and non-relaxing listening conditions. The intention was to select one track from within each genre (EDM, classical, and jazz) from within the low relaxation potential (non-relaxing listening condition) and high relaxation potential (relaxing listening condition) categories. To this end, each of the two tracks within the same genre under each relaxation potential category were compared, according to the criteria outlined in section *2.3. Pilot survey for the selection of unfamiliar music*. Bonferroni correction was applied to account for the seven comparisons made within each genre type in the sets of high and low relaxation potential tracks, resulting in an adjusted alpha of .007. See Supplementary Table 2 below for the means and standard deviations of the quality and familiarity ratings for each track.

With regard to the high relaxation potential tracks, paired samples t-tests showed there to be a significant difference between the electronic dance music (EDM) tracks *Escape (Fictivision Mix)* and *Untitled Instrumental No.4* in terms of valence only, *t*(16) = 3.12, *p* = .007, with the *Escape (Fictivision Mix)* track being rated with more positive valence. The classical music tracks *Carnival Overture (Op. 92)* and *Piano Concerto No. 1 in E-flat Major* showed significant differences in terms of energy, *t*(16) = 3.52, *p* = .003, and valence, *t*(16) = 3.12, *p* = .007, with the *Carnival Overture* track performing higher in both indices. Further, the jazz music tracks showed significant differences in terms of acousticness, *t*(16) = 3.23, *p* = .005, and valence, *t*(16) = 3.10, *p* = .007, with the track *Flik’s Machine* being rated with more positive valence and as being more acoustic than the *Quest for Coin* track. See Supplementary Figure 3 for visualisations of these comparisons.

Next, in terms of the low relaxation potential tracks, paired samples t-tests revealed significant differences between the EDM tracks *Ambient Track* and *Akiko* in energy, *t*(16) = 6.43, *p* < .001, valence, *t*(16) = 3.82, *p* = .002, and danceability, *t*(16) = 7.35, *p* < .001, with *Ambient Track* being considered as possessing low energy, neutral valence, and low danceability. Further, paired samples t-tests between the classical music tracks *Christmas Oratorio, BWV 248* and *Berlioz: Symphonie Fantastique* did not reveal any significant differences in terms of any of the track quality indices, however showed a tendency for the *Christmas Oratorio* track to be considered less energetic, neutrally valenced, more relaxing, and was liked more, which fulfilled our selection criteria to a greater extent than the *Symphonie Fantastique*. Finally, there was a significant difference between the jazz tracks *O Leazinho* and *Hello My Lovely* in terms of valence only, *t*(16) = 3.36, *p* = .004, with *O Leazinho* being rated as possessing more positive valence. See Supplementary Figure 4 for visualisations of these comparisons.

In a final check, we ran three additional paired-samples t-tests, this time comparing relaxingness ratings *across* conditions, in order to be sure that our chosen tracks within each genre indeed differed in terms of their relaxation potential. We found significant differences in levels of relaxingness between our EDM tracks, *Escape (Fictivision Mix)* and *Ambient Track*, *t*(16) = 7.17, *p* < 0.001, between our classical tracks, *Carnival Overture (Op. 92)* and *Berlioz: Symphonie Fantastique*, *t*(16) = 4.00, *p* = .001, as well as between our jazz tracks, *Flik’s Machine* and *Hello My Lovely*, *t*(16) = 3.87, *p* = .001.

**Supplementary Table 1.** Musical excerpts included in the pilot survey with identifying details.

| **Piece** | **Artist/Composer** | **Genre** | **Arousal** |
| --- | --- | --- | --- |
| Escape (Fictivision Mix)*^ | Fictivision | Electronic | High |
| Untitled Instrumental No.4 | Antoni Maiovvi | Electronic | High |
| Carnival Overture (Op. 92)*^ | Antonín Dvořák | Classical | High |
| Piano Concerto No. 1 in E-flat Major, S. 124 III Allegretto Vivace^ | Franz Liszt | Classical | High |
| The Flik Machine*^ | Randy Newman | Jazz | High |
| Quest for Coin | Ezra Collective | Jazz | High |
| Ambient Track* | MassiveMusic | Electronic | Low |
| Akiko^ | Guitar | Electronic | Low |
| Christmas Oratorio, BWV 248: Sinfonia in G*^ | Various Artists | Classical | Low |
| Berlioz: Symphonie Fantastique, Op. 14 – 3. Scene Aux Champs^ | Michael Tilson Thomas: San Francisco Symphony Orchestra | Classical | Low |
| O Leazinho^ | Brooks Williams | Jazz | Low |
| Hello My Lovely*^ | Enrico Pieranunzi | Jazz | Low |

Note. Those marked with * were the final tracks chosen for the study. In terms of tracks selected to be piloted, those marked with ^ were selected from Marti-Marca et al. (2020), and unmarked excerpts were selected by the experimenters independently.

**Supplementary Table 2.** Mean values (and standard deviations) of acoustic and aesthetic ratings of musical excerpts compared in a pilot survey for the purpose of selecting tracks to be used in the experimenter-selected conditions for the main experiment.

|  | **Genre** | **Energy** | **Acousticness** | **Valence** | **Danceability** | **Loudness** | **Relaxingness** | **Liking** | **Familiarity** |
| --- | --- | --- | --- | --- | --- | --- | --- | --- | --- |
| **Low Relaxation Potential** |  |  |  |  |  |  |  |  |  |
| Escape (Fictivision Mix) * | Electronic | 6.76 (0.44) | 1.47 (1.46) | 5.76 (1.30) | 6.00 (1.84) | 6.06 (1.03) | 1.59 (1.06) | 4.29 (2.08) | 1.06 (0.24) |
| Antoni Maiovvi - Untitled Instrumental No.4 | Electronic | 6.65 (0.61) | 1.35 (1.06) | 4.88 (0.86) | 5.35 (1.87) | 5.88 (1.17) | 1.53 (1.07) | 3.71 (1.83) | 1.12 (0.33) |
| Dvorak, Carnival Overture * | Classical | 6.29 (1.16) | 5.76 (2.08) | 5.29 (1.86) | 2.76 (2.19) | 5.82 (1.29) | 2.12 (1.45) | 4.00 (1.87) | 1.76 (0.90) |
| Liszt, Allegretto Vivace | Classical | 5.29 (1.21) | 5.65 (1.84) | 3.65 (1.90) | 1.88 (1.53) | 5.59 (1.54) | 1.94 (1.34) | 4.18 (1.94) | 1.53 (0.80) |
| Flik's Machine * | Jazz | 6.47 (0.72) | 5.76 (1.92) | 6.76 (0.44) | 5.82 (1.47) | 5.47 (1.07) | 3.35 (1.80) | 5.65 (1.54) | 1.41 (0.62) |
| Quest for Coin | Jazz | 6.12 (1.11) | 4.47 (1.70) | 6.12 (0.93) | 5.71 (1.76) | 5.41 (1.42) | 3.24 (1.44) | 5.24 (1.99) | 1.06 (0.24) |
| **High Relaxation Potential** |  |  |  |  |  |  |  |  |  |
| Ambient Track * | Electronic | 1.64 (1.17) | 3.47 (2.00) | 4.18 (1.78) | 1.24 (0.56) | 2.71 (1.49) | 5.47 (2.07) | 3.76 (1.95) | 1.12 (0.33) |
| Akiko | Electronic | 4.24 (1.03) | 3.82 (1.59) | 5.24 (1.09) | 3.76 (1.64) | 3.53 (1.12) | 4.82 (1.98) | 4.94 (1.89) | 1.12 (0.33) |
| Christmas Oratorio, BWV 248 * | Classical | 3.06 (1.14) | 6.12 (1.53) | 4.88 (1.45) | 3.06 (1.89) | 4.12 (1.76) | 5.24 (1.89) | 4.47 (1.81) | 1.47 (0.72) |
| Berlioz: Symphonie Fantastique | Classical | 3.41 (1.77) | 5.88 (1.58) | 4.65 (1.41) | 2.41 (1.73) | 4.18 (1.38) | 4.41 (2.03) | 4.00 (2.03) | 1.47 (0.51) |
| O Leazinho | Jazz | 4.06 (1.03) | 6.47 (0.94) | 5.59 (1.12) | 2.18 (1.07) | 2.94 (1.34) | 5.53 (1.84) | 5.00 (1.65) | 1.12 (0.33) |
| Hello My Lovely * | Jazz | 2.65 (1.46) | 5.29 (2.23) | 4.06 (1.48) | 2.59 (1.50) | 2.88 (1.45) | 5.41 (1.97) | 4.24 (1.92) | 1.18 (0.53) |

Note. N = 17. Excerpt pairs within each genre were compared using paired-samples t-tests for inclusion in the final set of unfamiliar tracks. Those marked with * were the final tracks chosen for the main experiment.

**Control subset analyses: cluster-based permutation analysis**

We addressed the potential confound of having large discrepancies between number of trials by running a control analysis, where the goal was to systematically minimise discrepancies as much as possible while retaining as many participants as we could. We conducted control analyses whereby we calculated the relative proportions of discrepancy between inter-condition trial numbers, then excluded participants with the highest levels of discrepancy. After participant exclusions, we reran the cluster-based permutation analysis within the alpha, gamma, theta, and delta bands along the main contrasts that were significant in the main analyses, with the following new trial numbers (M ± SD) and sample sizes [N]: (1) visual imagery (40.5 ± 31.9) vs. no visual imagery (20.4 ± 19.2) [20]; (2) spontaneous imagery (25.4 ± 21.5) vs. no visual imagery (20.4 ± 19.2) [20]; and (3) spontaneous imagery (25.0 ± 21.0) vs. deliberate imagery (17.3 ± 16.0) [20]. This subset analysis was not rerun on the deliberate imagery vs. no visual imagery contrast as no notable trial number discrepancies were found between them.

The analysis identified a just significant difference in alpha power suppression between **visual imagery and no visual imagery** probes within a comparable time range (–4.8 to –3.2 s, *p* = .050). Similar to the main analysis, there was alpha suppression between **spontaneous imagery and no imagery** probes in two clusters (cluster 1: –3.05 to –0 s, *p* = .016; cluster 2: –5.15 to –3 s, *p* = .030; see Supplementary Figure 1). In line with the main results, no clusters were identified in the alpha range between **deliberate and spontaneous imagery** probes (*p* = .281).

In contrast with the main analysis, there were no differences found in gamma power between **visual imagery and no imagery** probes (*p* = .148), or between **spontaneous imagery and no imagery** probes (*p* = .112). There was also no significant difference in gamma power between **deliberate imagery and spontaneous imagery** probes (*p* = .138). Finally, the analysis corroborated that there were no differences found between any of the contrasts with regard to the theta and delta bands.

These findings thus reduce concerns that key significant differences between states reported in the main findings are merely an artifact of discrepancies in trial numbers.

**Supplementary Figure 1.** Oscillatory differences between states (visual/spontaneous imagery and no visual imagery) as a function of time (s) relative to probe onset (0 s; indicated in the line plots by a dotted line along the y-axis) for alpha [8-13 Hz] band. (a/b) Time-frequency spectrograms averaged across channel sites of the power difference between visual/spontaneous imagery and no visual imagery probes. Clusters reflecting significant state differences are denoted by broken orange rectangles. (c/d) Alpha spectral power averaged over the channel sites of the clusters reflecting state differences between visual/spontaneous and no visual imagery states. The significant clusters are denoted by the black bars along the x-axis. (g) Topography of the clusters reflecting state differences between visual/spontaneous and no visual imagery states within the alpha band. Alpha clusters were segmented at 500 ms intervals (black markers denote electrodes that were present in the cluster). Rounded brackets surrounding sets of topoplots represent individual cluster groups.

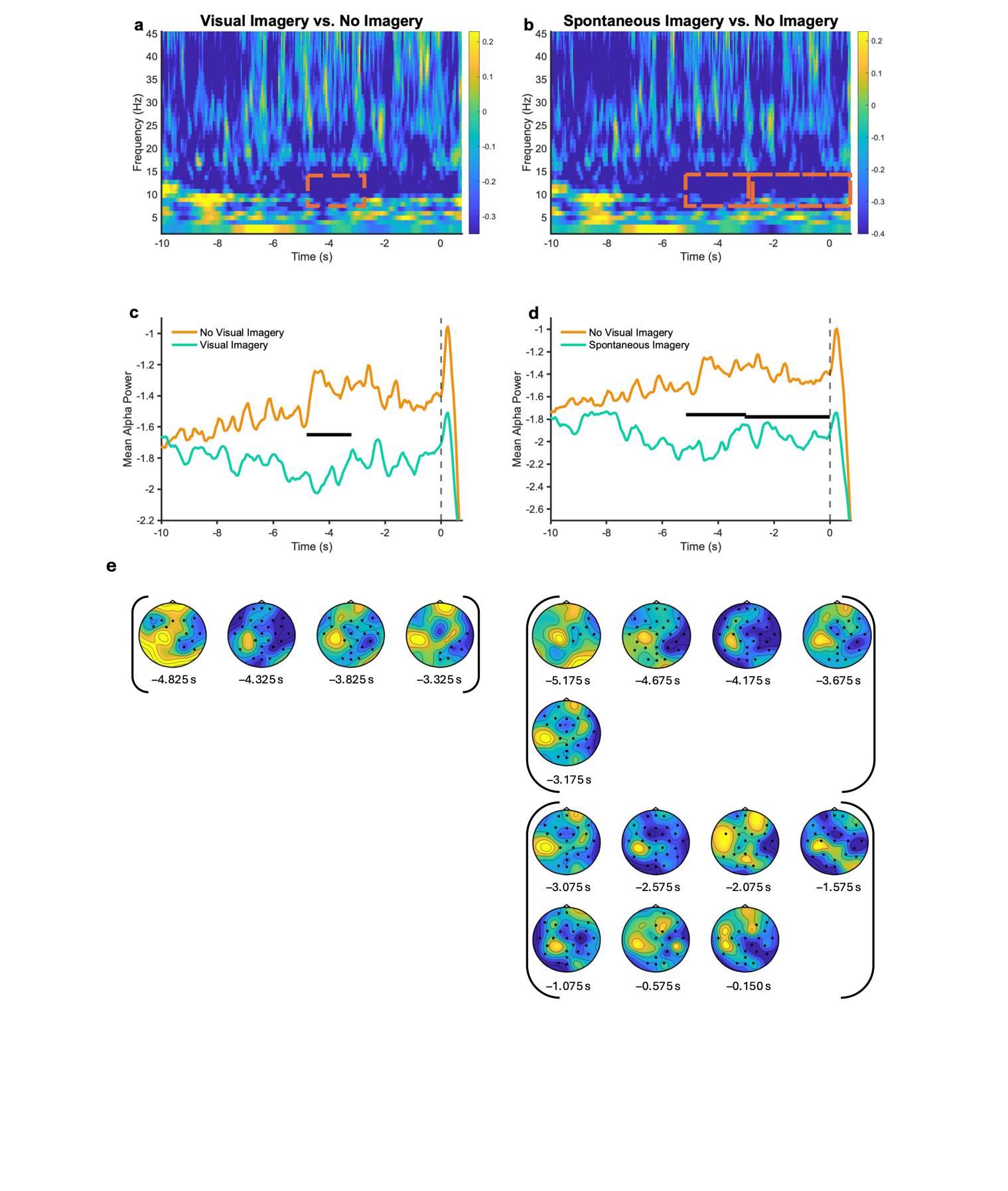

**Supplementary Figure 2.** Mean values (± standard error of the mean) of track quality responses for low relaxation potential tracks compared within genres (top plot: electronic dance music genre tracks, middle plot: classical music genre tracks, bottom plot: jazz music genre tracks). Asterisks indicate differences that reached corrected statistical significance (p < .007).

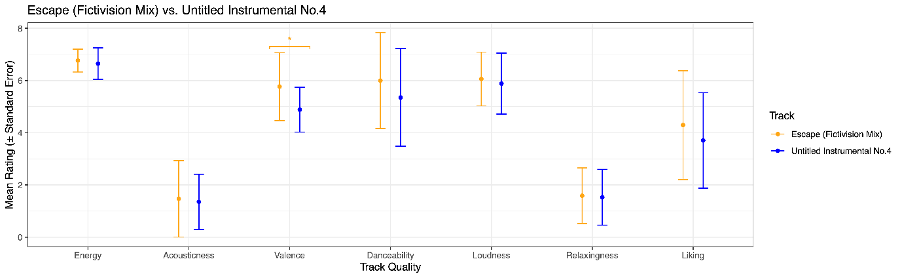

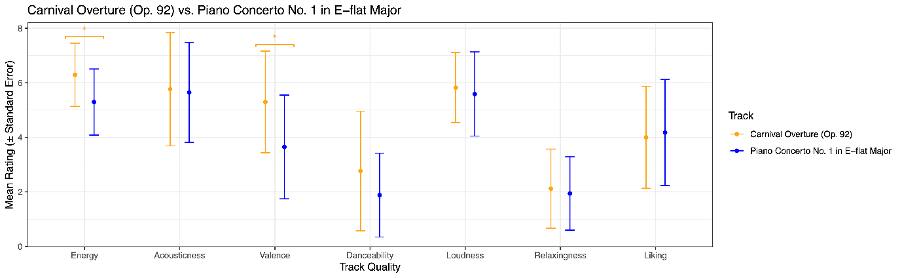

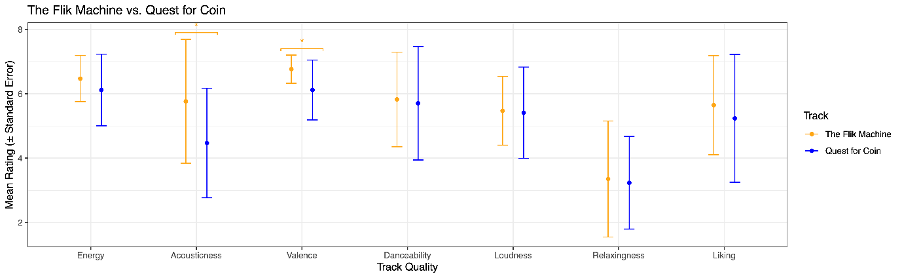

**Supplementary Figure 3.** Mean values (± standard error of the mean) of track quality responses for high relaxation potential tracks compared within genres (top plot: electronic dance music genre tracks, middle plot: classical music genre tracks, bottom plot: jazz music genre tracks). Asterisks indicate differences that reached corrected statistical significance (p < .007).

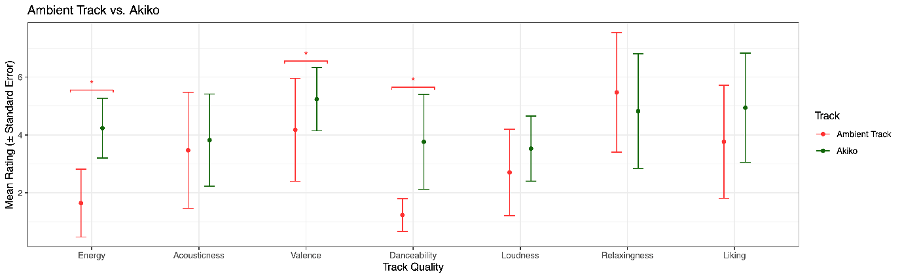

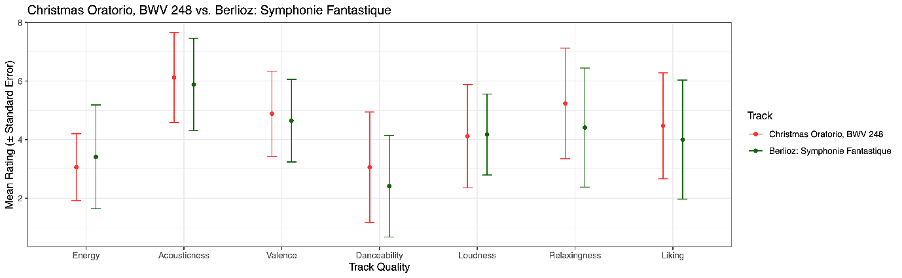

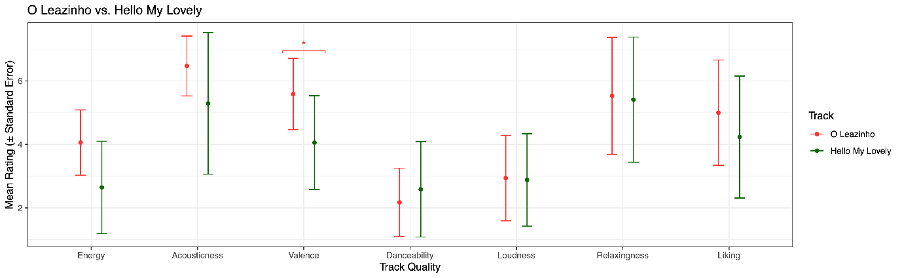

**Supplementary Figure 4.** Scatterplot and regression lines (with banded standard error of the mean) of relationships between rate of visual imagery probes and each of the individual difference indices (musical training, Psi-Q, and STOMP) between the two categories of listening conditions: those with high or low Relaxation Potential (left) and high and low Familiarity (right).

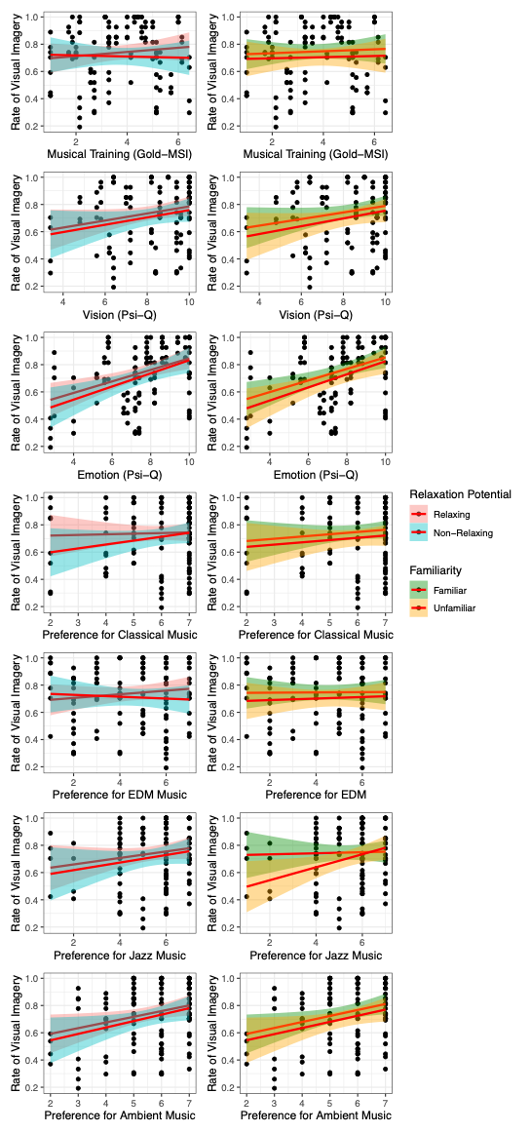

**Supplementary Figure 5.** Scatterplot and regression lines (with banded standard error of the mean) of relationships between rate of spontaneous visual imagery probes and each of the individual difference indices (musical training, Psi-Q, and STOMP) between the two categories of listening conditions: those with high or low Relaxation Potential (left) and high and low Familiarity (right).

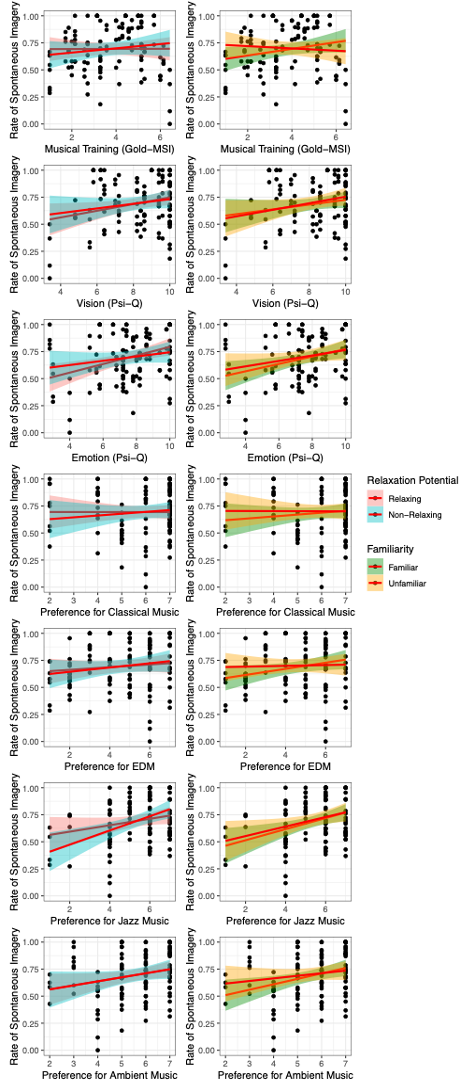
